# Atribacteria reproducing over millions of years in the Atlantic abyssal subseafloor

**DOI:** 10.1101/2020.07.10.198200

**Authors:** Aurèle Vuillemin, Sergio Vargas, Ömer K. Coskun, Robert Pockalny, Richard W. Murray, David C. Smith, Steven D’Hondt, William D. Orsi

## Abstract

How microbial metabolism is translated into cellular reproduction under energy-limited settings below the seafloor over long timescales is poorly understood. Here, we show that microbial abundance increases an order of magnitude over a five million-year-long sequence in anoxic subseafloor clay of the abyssal North Atlantic Ocean. This increase in biomass correlated with an increased number of transcribed protein-encoding genes that included those involved in cytokinesis, demonstrating that active microbial reproduction outpaces cell death in these ancient sediments. Metagenomes, metatranscriptomes, and 16S rRNA gene sequencing all show that the actively reproducing community was dominated by the candidate Phylum “*Candidatus* Atribacteria”, which exhibited patterns of gene expression consistent with a fermentative, and potentially acetogenic metabolism. “*Ca.* Atribacteria” dominated throughout the entire eight million-year-old cored sequence, despite the detection limit for gene expression being reached in five million-year-old sediments. The subseafloor reproducing “*Ca.* Atribacteria” also expressed genes encoding a bacterial micro-compartment that has potential to assist in secondary fermentation by recycling aldehydes and, thereby, harness additional power to reduce ferredoxin and NAD^+^. Expression of genes encoding the Rnf complex for generation of chemiosmotic ATP synthesis were also detected from the subseafloor “*Ca*. Atribacteria”, as well as the Wood-Ljungdahl pathway that could potentially have an anabolic or catabolic function. The correlation of this metabolism with cytokinesis gene expression and a net increase in biomass over the million-year-old sampled interval indicates that the “*Ca*. Atribacteria” can perform the necessary catabolic and anabolic functions necessary for cellular reproduction, even under energy limitation in millions of years old anoxic sediments.

**Importance:** The deep subseafloor sedimentary biosphere is one of the largest ecosystems on Earth, where microbes subsist under energy-limited conditions over long timescales. It remains poorly understood how mechanisms of microbial metabolism promote increased fitness in these settings. We discovered that the candidate bacterial Phylum “*Candidatus* Atribacteria” dominated a deep-sea subseafloor ecosystem, where it exhibited increased transcription of genes associated with acetogenic fermentation and reproduction in million-year old sediment. We attribute its improved fitness after burial in the seabed to its capabilities to derive energy from increasingly oxidized metabolites via a bacterial micro-compartment and utilize a potentially reversible Wood-Ljungdahl pathway to help meet anabolic and catabolic requirements for growth. Our findings show that “*Ca*. Atribacteria” can perform all the necessary catabolic and anabolic functions necessary for cellular reproduction, even under energy limitation in anoxic sediments that are millions of years old.

## Introduction

Marine sediments contain an ubiquitous “deep biosphere” (1) extending at least as far as 2,500 meters below the seafloor (mbsf) (2), which consist of active and dormant cells (3–5) with measurable impacts on subseafloor biogeochemical processes (6). At abyssal water depths in the deep sea, the subseafloor communities generally are less sampled (7) compared to those in continental shelf sediments that have higher activities and rates of microbial sulfate reduction (8–10). At abyssal depths under the oligotrophic ocean gyres, the sedimentation rates are low, ranging from 1 to 5 meters of sediment deposited per million years (11, 12). These abyssal subseafloor communities have extremely low metabolic activity (13) and live near the energy limit to life (6, 14). As a result of this, the deep biosphere of marine sediment is generally characterized by net death, and it remains poorly understood to what extent microbial activity is translated into cellular reproduction in the deep subseafloor (15, 16).

Many subseafloor microbes exhibit viability since they actively take up carbon and nitrogen in incubation experiments (4, 17), indicating potential for microbial growth in energy-limited anoxic subseafloor sediments. Microbial activities can also be stimulated at redox interfaces deep below the seafloor over geological timescales (3, 18). However, the capacity of microbes to reproduce in abyssal subseafloor ecosystems close to the energy limit to life (6, 14) is particularly unconstrained, given the extreme scarcity of organic substrates in these settings (19). There is reason to suspect that cellular reproduction in the abyssal subseafloor is minimal since microbial biomass tends to decrease an order of magnitude over the top 10 m of sediment in all abyssal locations yet sampled, reaching the detection limit for life at relatively shallow subseafloor depths of ca. 15 mbsf (12). This follows the global trend whereby subseafloor microbes tend to die faster than they grow (1), particularly in the top 10 m of marine sediment.

Here, we report an exception to this global trend in anoxic deep-sea clay recovered from an abyssal water depth of >5,500 m in the North Atlantic, characterized by an ultra-slow sedimentation rate of ca. 3 m per million years. In contrast to oxic abyssal red clay where microbial abundance decreases several orders of magnitude over the top 10 mbsf (12, 20), we show here that microbial abundance in the anoxic abyssal clay increases an order of magnitude from the seafloor down to 15 mbsf (spanning ca. 5 million years). We then proceeded to use metatranscriptomics to further investigate the anaerobic metabolic mechanisms that explain this net growth in the size of the subseafloor microbial ecosystem over multimillion year timescales.

## Results and Discussion

### Sediment biogeochemistry

We obtained deep-sea clay sediment from 5,515 meter water depth in the ultra-oligotrophic open ocean of the North Atlantic. This coring site (KN223-15) is characterized by a mean sedimentation rate of ca. 3 m per million years (11). Samples ranged from 0.1 to 30 meters below seafloor (mbsf). Given the mean sedimentation rate, the deepest sample has an approximate age of 9 to 10 million years. Oxygen and nitrate penetration into the sediment are restricted to the top mm of sediment, as they were detectable in the bottom water but below detection in the uppermost portion of the core at 0.02 and 0.03 mbsf, respectively (Fig. S1). The abyssal sediments of the North Atlantic are typically oxic red clay that tend to have O_2_ penetrating many meters into the seafloor (11, 21), but the subseafloor microbial ecosystem sampled here is unique in the sense that the sediments are anoxic despite having ultra-slow sedimentation rates. Moreover, while the sediment of our abyssal subseafloor core displays sulfate (SO_4_^2−^) concentrations about 29 mM at 0.02 mbsf and is fully anoxic downward, the rates of anaerobic microbial SO_4_^2−^ reduction over the top 10 meters of sampled sediment are low (−3.8 × 10^−4^ mol SO_4_^2−^ × m^3^ yr^−1^) and below detection underneath, resulting in pore water SO_4_^2−^ remaining over 20 mM throughout the entire core (Fig. 1A). This profile is very similar to profiles observed previously in anoxic sediments from other oligotrophic regions, such as the Eastern Equatorial Pacific, where sediments are anoxic but community metabolic activity is too slow to consume all of the available SO_4_^2−^ (6, 22).

**Figure 1.**
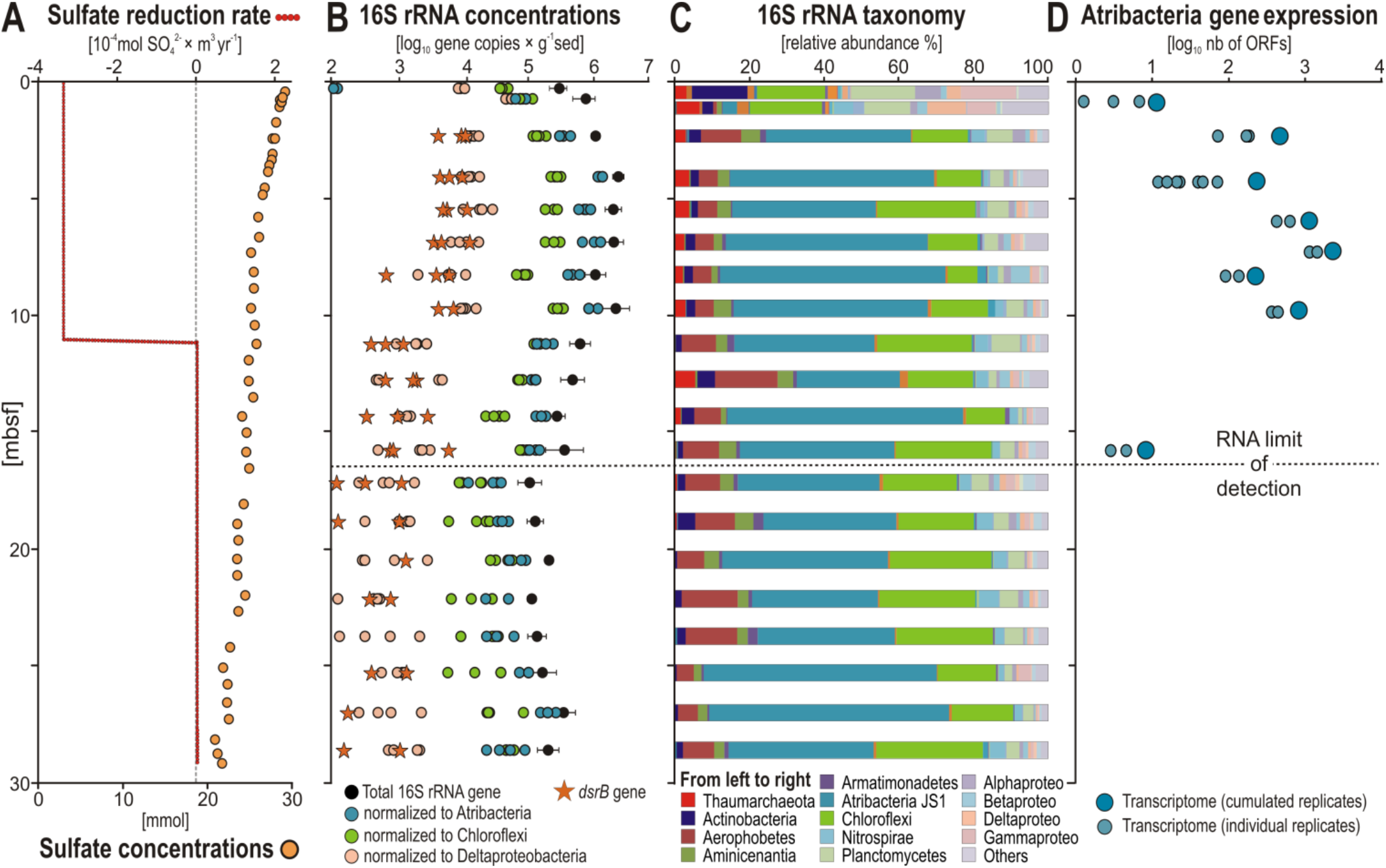
Biogeochemistry, microbial diversity and abundance in the anoxic sediment of North Atlantic Site KN223-15. (**A**) Profiles of mean net SO_4_^2−^ reduction rates and dissolved SO_4_^2−^ concentrations. (**B**) Quantitative polymerase chain reaction (qPCR) of 16S rRNA (black dots) and *dsrB* (orange stars) genes, and qPCR-normalized abundance of Atribacteria (blue), Chloroflexi (green) and Deltaproteobacteria (light orange) 16S rRNA genes. Error bars correspond to standard deviation (2 σ) based on four biological replicates. (**C**) Diversity of 16S rRNA genes based on three to four biological replicates. **(D**) Number of ORFs attributed to Atribacteria in the metatranscriptomes as a function of depth. Light blue circles are the number of ORFs detected in individual metatranscriptome replicates, and the larger dark blue circles are the total number of unique ORFs detected when summing across all replicates. The top two samples in panels B and C (0.1 – 0.3 mbsf) were recovered by gravity coring, the deeper samples (0.5 – 29 mbsf) were recovered via long piston coring. Note in panels B and C that an increase in abundance of Atribacteria in the upper 1 mbsf coincides with their higher level of gene expression in panel D.

### Abundance and diversity of the microbial communities

The density of 16S rRNA gene copies per gram of wet sediment exhibits a subsurface peak in abundance, increasing one order of magnitude (2.5 × 10^5^ to 2.3 × 10^6^ copies) from 0.2 to 3.5 mbsf and remaining >10^6^ between 3 and 10 mbsf. Thereafter, microbial 16S rRNA gene abundances decrease gradually to a minimum of 9.9 × 10^4^ copies at 17 mbsf (Fig. 1B). In the uppermost sediment samples (0.1 - 0.4 mbsf), Actinobacteria, Planctomycetes, Delta- and Gammaproteobacteria dominate the community (Fig. 1C). Below this depth starting at 0.5 mbsf, the relative abundance of “*Ca*. Atribacteria” rapidly increase from <5 % to >40 % (Figs. 1B-1C) in all four biological replicates sampled (Fig. S2, S3). The “*Ca*. Atribacteria” is only represented by 3 OTUs, with one single OTU accounting for up to 40 % of the whole community throughout the record (Fig. 2A-B) which was consistent across three to four biological replicates (Figs. S2). This dominant OTU (Fig. 2A) was most closely affiliated with unpublished 16S rRNA gene sequences from deep subseafloor sediments from the Nankai Trough (23) and a potential gas hydrate region of southwest Taiwan subseafloor sediments (Fig. 2B).

**Figure 2.**
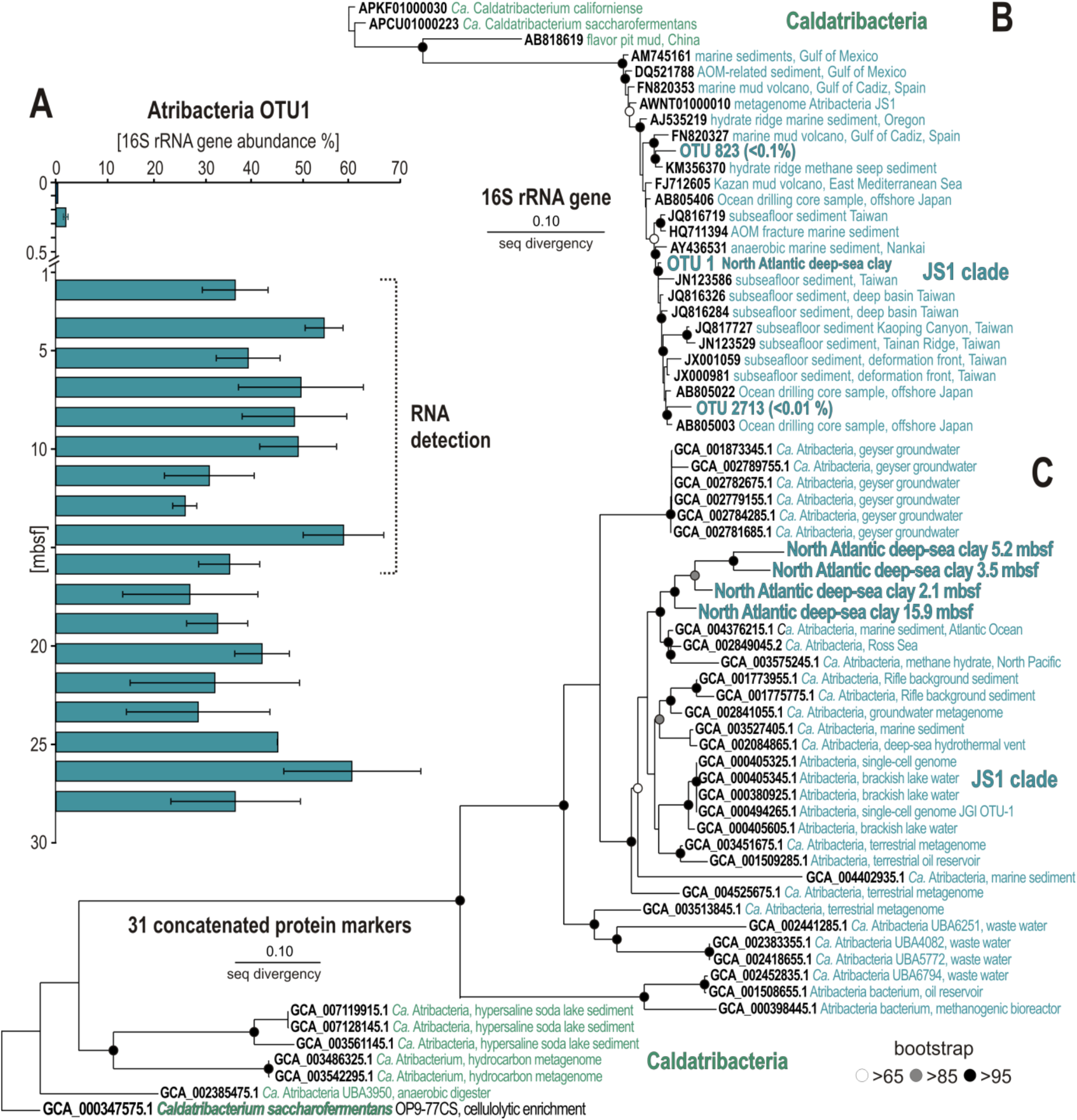
Relative abundance and phylogenetic analyses of the most abundant OTU (“ *Ca*. Atribacteria”) and its phylogenomic protein markers. (**A**) The histogram shows the relative abundance of the most abundant OTU (“*Ca*. Atribacteria”), error bars are standard deviations across 3 to 4 biological replicates (Fig. S2). (**B**) Phylogenetic analysis of 16S rRNA genes (V4 hypervariable region) showing the affiliation of all three “*Ca*. Atribacteria” OTUs to the JS1 clade. (**C**) Phylogenetic tree based on 31 concatenated protein markers identified with AMPHORA2 (33). The monophyletic clade including our samples in the tree have >95% similarity to the sister clade with taxa from the Atlantic Ocean, Ross Sea, and North Pacific methane hydrate subseafloor sediment.

Following a protocol for quantitative normalization of barcoded 16S gene sequence data (24), we normalized group-specific 16S rRNA gene abundances by dividing the total number of 16S rRNA gene copies determined via quantitative PCR (qPCR) by their affiliated 16S rRNA gene relative abundances (fractional percentage of total sequences per sample). This showed that “*Ca*. Atribacteria” dominates the community (Figs. 1B), which is attributed to a single 16S OTU (Fig 2). We considered the potential influence of multiple copies of 16S rRNA genes (25) on the result and searched for all “*Ca*. Atribacteria” genomes in the JGI database sequenced to date, which revealed that all sequenced genomes from this group have only one 16S rRNA gene copy, whereas Deltaproteobacteria and Chloroflexi have a median of two 16S rRNA gene copies (up to four) per genome (25). Thus, despite having an average lower 16S rRNA gene copy number compared to Chloroflexi and Deltaproteobacteria, “*Ca*. Atribacteria” relative abundance was high in our 16S rRNA gene sequence dataset, even suggesting that our data actually underestimates the abundance of “*Ca*. Atribacteria”. These profiles show exponential net increases of “*Ca*. Atribacteria” and Chloroflexi within the upper 10 m of sediment, whereas those of Deltaproteobacteria increase at 0.4 mbsf, but recede rapidly over time from 10^4^ to 10^3^ copies or less, reaching our limit of detection (10^2^ gene copies × gram^−1^ wet sed) at the bottom of the core. The vertical profile of the beta subunit of the dissimilatory sulfate reductase (*dsrB*) gene runs parallel to that of Deltaproteobacteria 16S rRNA genes, which points to Deltaproteobacteria as the main sulfate-reducing bacteria (SRB) in this anoxic sediment. This inference is supported by the abrupt 10-fold drop in *dsrB* and Deltraproteobacteria 16S rRNA gene sequences at 10 mbsf (Fig. 1B), the same depth at which net SO_4_^2−^ reduction decreases to below detection levels (Fig. 1A). The abundance of SRB is lower compared to organic rich shelf sediments (26, 27), which may be related to the relatively low sedimentation rate at our sampling location since SO_4_^2−^ reduction rates are correlated to the square of sedimentation rate (9). The relative slowness of SO_4_^2−^ reduction at our sampling location is readily evident in the profile of dissolved SO_4_^2−^ concentrations, which remain above 20 mM throughout the entire core (Fig. 1A).

### Metagenomic analysis

Metagenomes from five depths were sequenced at an average depth of 8.4 million reads (± 2.5 millions) (Table S3). De novo ‘binning’ based methods for creating metagenome-assembled genomes (MAGs) are useful for discovering new taxa when distantly related genomes in databases preclude similarity-based searches (28–31). Because of the ultra-low DNA concentrations extractable from these abyssal clay sediments, we were only able to obtain metagenomic data after amplifying the extracted DNA using multiple displacement amplification (MDA). Our attempts at *de novo* binning of the MDA products revealed a selective amplification of short fragments that precluded binning and completion of high-quality MAGs (Table S2). Specifically, manual curation of Maxbin results using Anvi’o (32) produced 12 bins at relatively low levels of genome completeness (17-44%) that could be assigned to “*Ca*. Atribacteria” (Fig. S4). Thus, in order to increase the annotation of putative functions of open reading frames (ORFs) in metagenomes from the “Ca. Atribacteria” that could not be assembled into bins we also applied a bioinformatics pipeline, whereby ORFs encoded on *de novo* assembled contigs were searched for similarity against a large aggregated genome database of predicted proteins from all Atribacteria MAGs and single cell genomes (SAGs) sequenced to date, including all published data from subsurface metagenomics studies (see Materials and Methods). We then extracted all the ORFs having a predicted protein from a “*Ca*. Atribacteria” genome as a top BLASTp hit and ran a phylogenomic analysis based on 31 phylogenomic markers using AMPHORA (33). This phylogenomic analysis demonstrates that our previously published method (34, 35), can recover ORFs with high similarity (>95% amino acid similarity) to predicted proteins in previously sequenced “*Ca*. Atribacteria” genomes (Fig. 2C). The phylogenomic analysis shows that the atribacterial ORFs from all sampled depths form a monophyletic clade sister to those from deep-sea sediments of the Atlantic Ocean, Ross Sea, and Pacific Ocean methane hydrates within the JS1 clade (Fig. 2C). This high level of similarity to existing “*Ca*. Atribacteria” genomes enabled similarity-based assignment of “*Ca*. Atribacteria” ORFs in our samples (Fig. S5). As further evidence of this, the median similarity of ORFs to their top hit in the database in all metatranscriptome and metagenome datasets >60% indicates that the ORFs in our samples had relatively high similarity to existing predicted proteins in the database (Fig. S6).

We confirmed the accuracy of this previously published approach (34, 35) by an *in silico* test for true and false positive annotations based on 151 randomly selected peptide fragments extracted from “*Ca*. Atribacteria” predicted proteomes, as well as other bacterial and archaeal genomes (Fig. S7). In this analysis, 50 randomly selected predicted proteins were randomly cut into peptide fragments ranging from 20 – 140 amino acid residues in length, in order to replicate partial ORFs typically recovered in metagenomes. The random peptide fragments were then searched against the large aggregate database (34, 35) for their top hits with BLASTp. This *in silico* test showed that 100% of all randomly cut peptide fragments from Atribacteria were true positives, they had the same Atribacteria genome as a top BLASTp hit. This shows that our use of a similarity-based approach for ORF annotations (34–36) is adequate for assigning ORFs encoded on *de novo* assembled contigs to groups at high taxonomic levels including those derived from the Candidate Phylum Atribacteria.

### Gene expression analysis

Metatranscriptomes from eight depths were produced in biological replicates (two to seven replicates per depth: Fig. 1D) and sequenced at an average depth of 4.0 million reads (± 1.5 million) with 91,199 contigs across all samples sequenced (Table S3). Similar to the 16S rRNA gene abundances, there was a subsurface peak in the number of unique expressed ORFs assigned to “*Ca*. Atribacteria” that increases exponentially between 3.5 and 10 mbsf, which was consistent across multiple replicate metatranscriptomes from each depth, and at 16 m our RNA limit of detection was reached (Fig. 1D). We defined this depth as our RNA detection limit, because the number of unique ORFs assigned to ‘*Ca*. Atribacteria’ were no longer detectable and below this depth the only ORFs annotated were those assigned to groups of known contaminants from molecular kits including those from human skin and soil (37). Many of these same group are common laboratory contaminants found in dust samples from our lab in 16S rRNA gene surveys (38), and include *Pseudomonas*, *Rhizobium*, *Acinetobacter*, and *Staphylococcus*. We interpreted the sudden dominance of ORFs with similarity these common contaminants below 15.9 mbsf, and the lack of ORFs annotated to the ‘*Ca*. Atribacteria’ that dominate throughout the core (Fig 1B), to be indicative of our limit of RNA detection. Presumably, this limit is reached because a lower amount of extracted RNA from the *in situ* active community becomes overprinted by background “noise” from contaminating DNA (or RNA) derived from the kits, aerosols, or other laboratory contaminants.

There was a statistically significant correlation (r-values = 0.59, 0.61; p-values = 0.016, 0.014) between the abundance of Atribacteria 16S rRNA genes and expressed ORFs (Table S1). The number of expressed ORFs correlated with 16S rRNA gene quantities, which was consistent between the entire dataset (total bacteria) and when comparing number of ORFs expressed per group to the qPCR normalized abundance of the same taxonomic group for the four group with the highest number of detected ORFs in the metatranscriptomes (Atribacteria, Deltaproteobacteria, Chloroflexi, and Archaea) (Figs. 3A and S8).

**Figure 3.**
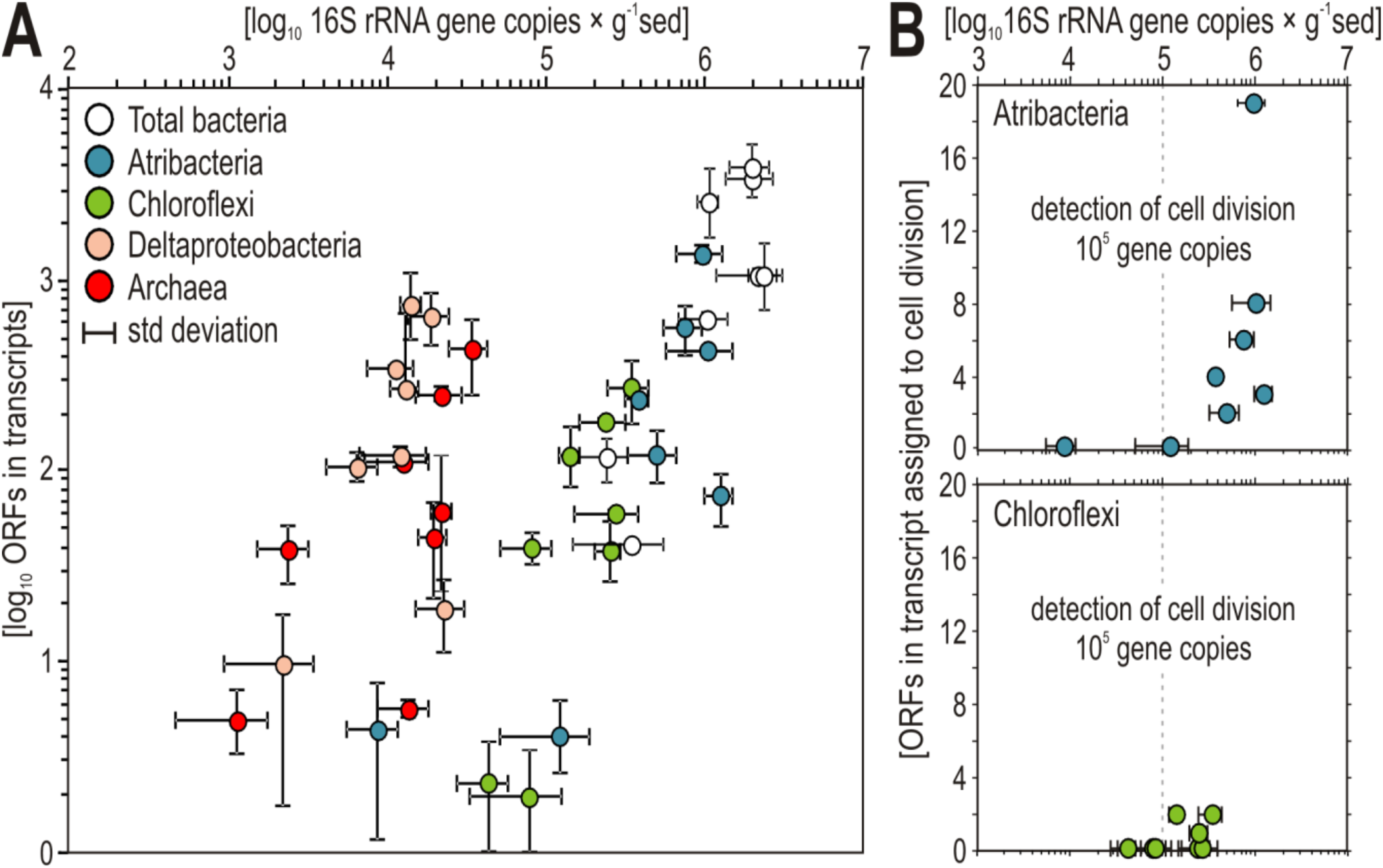
Group specific transcriptional activity correlates with abundance. **(A)** Abundance of 16S rRNA genes (X axis) normalized to total bacteria (white), Atribacteria (blue), Chloroflexi (green), Deltaproteobacteria (light orange) and Archaea (red) 16S rRNA sequences plotted against their respective number of ORFs in the assigned to the same groups in the metatranscriptomes (Y axis). Error bars represent standard deviation across biological replicates (in the case of three or more) or ranges (in the case of two replicates). **(B)** Abundance of 16S rRNA genes normalized to “*Ca*. Atribacteria” (blue) and Chloroflexi (green) plotted against their respective number of ORFs in the transcripts assigned to cell division, showing that cell division is only detectable at 16S rRNA gene densities > 10^5^ copies.

### Evidence for subseafloor reproduction

The steadily increasing abundance of “*Ca*. Atribacteria” over time (since sediment deposition) are apparently due to cells undergoing cytokinesis, as the 16S rRNA gene abundances are correlated with transcription of genes encoding proteins involved in cell division and cell shape determination such as FtsAEKQWZ, MreBC, and RodA (Figs. 3B and S9) (39, 40). The expressed genes encoding the Fts proteins form the “divisome” (41), which includes the FtsZ ring, a cytokinetic protein ring that localizes at the cell division site prior to cytokinesis in dividing bacterial cells and pumps the replicated chromosome into the daughter cell (42). These genes are expressed during cellular division in “*Ca*. Atribacteria”, indicating exponential growth phase (43). Thus, while we do not report measurements of biomass turnover such as amino acid racemization (44, 45) or stable isotope probing (4, 17), the steadily increasing microbial biomass of “*Ca*. Atribacteria” with increasing sediment depth together with the detected correlation between their abundance and transcriptional activity (Fig. 3A), and the transcription of ORFs encoding FtsZ ring proteins (Figs. 3B and S9), strongly suggest that the higher abundances of “*Ca*. Atribacteria” between 0.5 and 10 mbsf are due to actively dividing cells. In this zone of apparent increased reproduction, the number of unique expressed ORFs correlated significantly with the number of 16S rRNA gene copies from the dominant groups (Figs. 1 and 3). The most parsimonious explanation for this is that more expressed ORFs from Atribacteria are detected in these depths because there are higher numbers of metabolically active Atribacteria producing mRNA transcripts.

The sediment ages in this interval spans several million years, and thus this higher number of metabolically active microbes originated from “*Ca*. Atribacteria” that slowly reproduced through binary fission and cytokinesis over multimillion year timescales. Because genes involved in formation of the divisome, FstZ ring, and cytokinesis were only expressed in depths where the highest 16S rRNA gene copies were found (Fig. 4B), the higher number of 16S rRNA gene copies at those depths is likely due to higher numbers of actively reproducing atribacterial cells. Active cells have higher numbers of ribosomes, and thus higher copies of 16S rRNA per cell (46), but our qPCR assay targeted DNA (the bacterial chromosome), not RNA (expressed genes), thus discarding active rRNA synthesis as a confounding factor in our bacterial abundance estimations. Since “*Ca*. Atribacteria” has only one 16S rRNA gene copy, it is possible to conclude that the observed ten-fold increase in atribacterial 16S rRNA gene copy numbers over the top 10 mbsf requires chromosomal (genome) replication which, in bacteria, occurs in actively dividing cells prior to cytokinesis and binary fission (46), assuming steady state microbial input in sediment over millions of years. The only alternative explanation to our observations would require an increase in 16S rRNA copy number in the chromosome of “*Ca*. Atribacteria” over the top 10 mbsf followed by its subsequent decrease in the same chromosome below 10 mbsf. Such an incredibly fast rate of genome evolution affecting the highly conserved 16S gene and then acting in a reversible way after 10 mbsf is inconceivable, especially considering this gene evolves at a rate of 1% every 100 million years (47) and our deepest sampled depth is roughly 10 million years old. Thus, the most likely explanation is that the detection of more 16S rRNA gene copies from the dominant OTU of “*Ca*. Atribacteria” (Fig. 2A) is due to more cells, each containing one chromosome with one copy of the 16S rRNA gene. The correlation of these 16S rRNA genes with higher numbers of expressed genes from “*Ca*. Atribacteria” over the top 15 mbsf (Figs. 1D, and 3) can be attributed to a single clade (Fig. 2) that has been slowly reproducing and increasing in abundance over millions of years.

**Figure 4.**
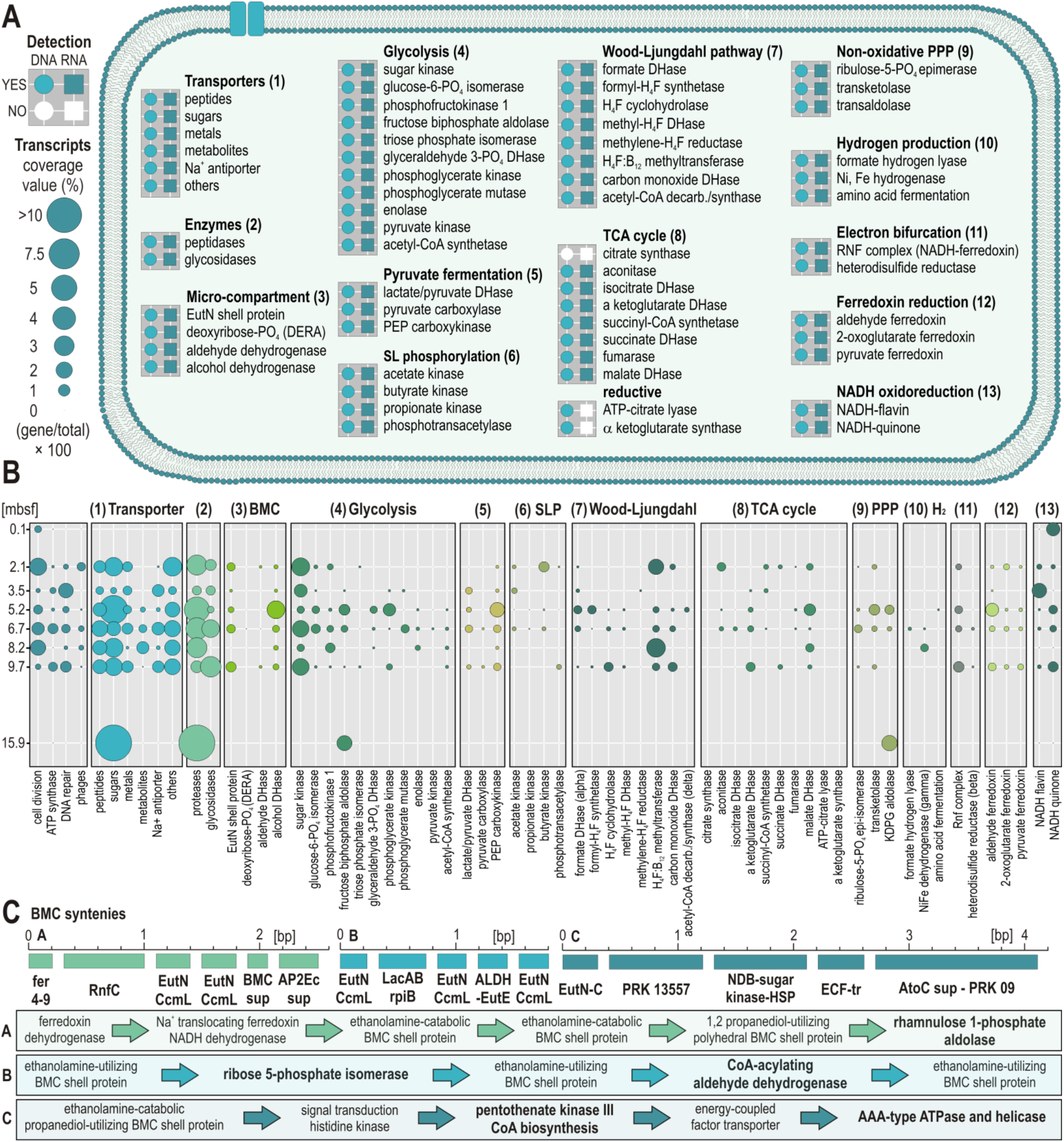
Metabolic potential and transcriptional activity of “*Ca*. Atribacteria” and its micro-compartment. (**A**) Presence (colored) or absence (white) of ORFs assigned to “*Ca*. Atribacteria” encoding predicted proteins in metagenomes (circles) and metatranscriptomes (squares). (**B**) Bubble plots showing the coverage values [% total reads] of expressed genes identified for “*Ca*. Atribacteria” at eight different depths. The numbering of the groups in panel A corresponds to the same numbering in panel B. DERA: 2-deoxy-D-ribose 5-phosphate aldolase. (**C**) Sequences of gene synteny related to the BMC present in the cell of “*Ca*. Atribacteria”. These three syntenies provide evidence for metabolic use of aldehydes and alcohols via dehydrogenases with biosynthesis of acetyl-coenzyme A and regeneration of NAD^+^.

### Predicted metabolism for “Ca. Atribacteria”

Our finding that “*Ca*. Atribacteria” expressed the highest number of protein-encoding genes compared to all other groups (Fig. 3) is furthermore consistent with their high levels of gene transcription in deep subseafloor sediments of the Baltic Sea (48). Below 15 mbsf, the microbial ecosystem transitions to net death since microbial abundances decrease by two orders of magnitude, RNA levels decline to below detection, and the net SO_4_^2−^ reduction rate is below our detection limit (Fig. 1). However, “*Ca*. Atribacteria” remains the dominant group throughout the core, even below 10 mbsf. Its continued dominance is consistent with its abundance in other deep subseafloor settings (49–52), and the findings that dominant taxa in the subseafloor community do not necessarily require cellular reproduction to outcompete other taxa, but can reach higher relative abundances from lower mortality compared to their competitors over long timescales (53, 54). Similar to what has been predicted from genomic studies (55–58), the gene transcription data strongly indicate that the “*Ca*. Atribacteria” dominating throughout the core actively utilizes a sugar-based acetogenic metabolism (Fig. 4A). This is also consistent with gene transcription from “*Ca*. Atribacteria” in deep subseafloor Baltic Sea sediments that showed active utilization of trehalose (48).

The Wood-Ljungdahl pathway (WLP) can be used catabolically to achieve redox balance and regenerate NAD^+^ and oxidized ferredoxin to thereby increase anaerobic metabolic efficiency (59). In acetogenic bacteria, this is coupled to the Rnf complex at the membrane that utilizes either ferredoxin or NAD^+^ as the terminal electron acceptor, driving a Na^+^ ion gradient and ATP synthesis (Fig. 5) (60). Several lines of evidence in our gene transcription data indicate that “*Ca*. Atribacteria” utilizes a similar metabolism.

**Figure 5.**
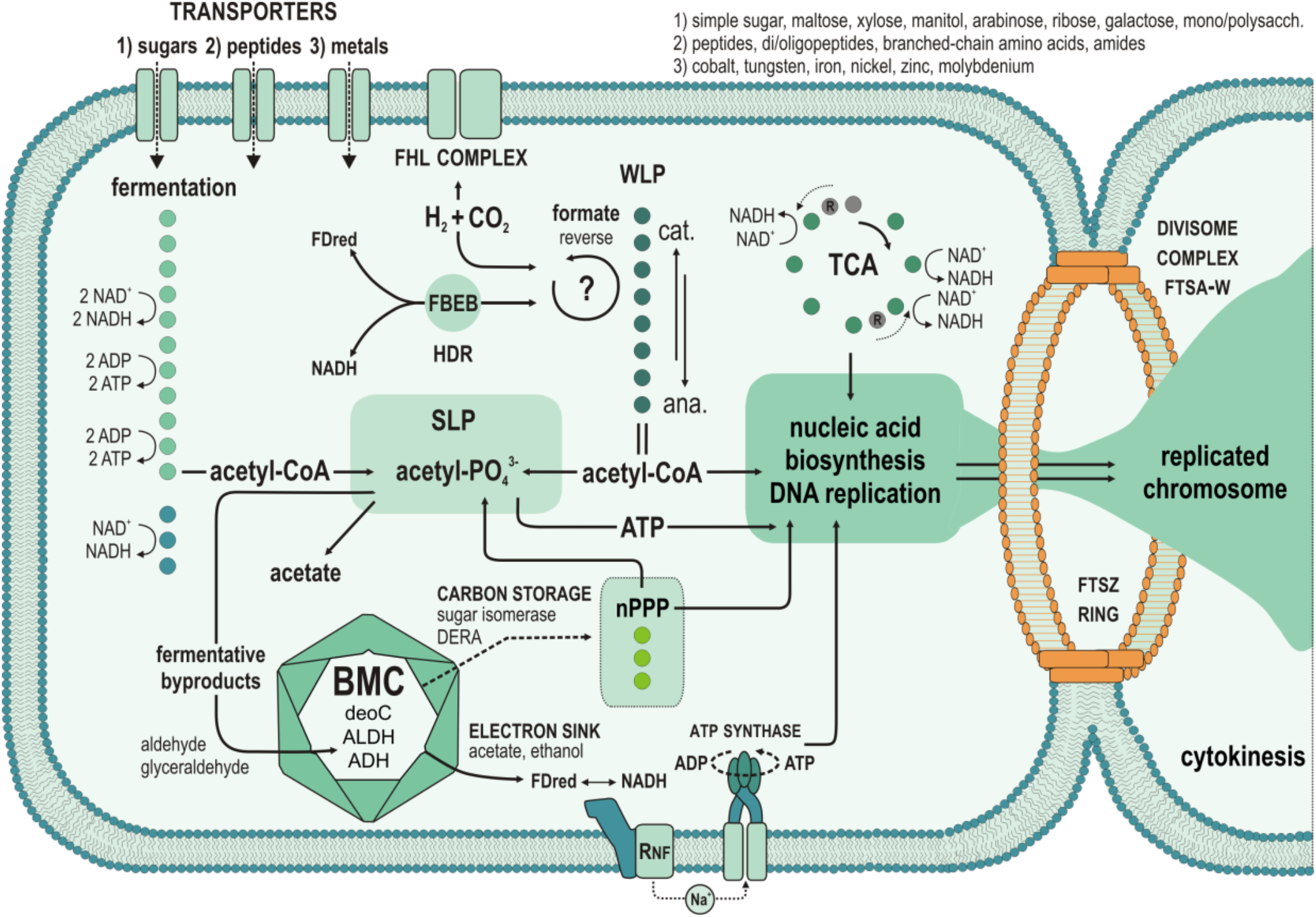
Potential “*Ca*. Atribacteria” physiology based on metagenomes and transcriptomes. The figure is based on the known interlinking of energy metabolism pathways for (homo)acetogenic bacteria (59) and BMC metabolic functions (57), as we all the function of the FstZ ring and divisome (41). Colored dots correspond to the successive enzymatic steps (as listed in Fig. 4), e.g. fermentation corresponds to glycolysis (11 dots) and lactate fermentation (3 dots). Actively expressed genes encoding the divisome complex (i.e. FtsAEKQWZ) are also depicted in Fig. S9. nPPP: non-oxidative pentose phosphate pathway. SLP: substrate level phosphorylation, DERA: 2-deoxy-D-ribose 5-phosphate aldolase, ALDH: aldehyde dehydrogenase, ADH: alcohol dehydrogenase, FBEB: flavin-based electron bifurcation, FHL: formate hydrogen lyase, WLP: Wood-Ljungdahl pathway.

Specifically, “*Ca*. Atribacteria” expressed transcripts encoding ORFs with similarity to proteins involved in sugar fermentation and the WLP (Fig. 4B). Genes encoding enzymes of the WLP were all present (Fig. 4A) and differently expressed by “*Ca*. Atribacteria”, as well as those encoding proteins involved in glycolysis, fermentation, and electron bifurcation (Fig. 4B). Transcription of ORFs by “*Ca*. Atribacteria” with similarity to enzymes known to be involved in electron bifurcation that results in production of molecular hydrogen included the beta subunit of the heterodisulfide reductase, formate hydrogen lyase (61), gamma subunit of the NiFe hydrogenase (62), and subunit alpha of the reversible formate dehydrogenase (63). The formate hydrogen lyase complex combines NiFe hydrogenase and soluble formate dehydrogenase to couple formate and/or CO-dependent hydrogen production to the generation of Na^+^ motive force to generate ATP. This enzymatic complex allows growth on formate disproportionation (64), during which two electrons are passed to hydrogenase and two protons are neutralized (65). The gene expression data indicate that the ‘Ca. Atribacteria’ produce the chemiosmotic Na^+^ gradient using the Rnf complex for ATP synthesis (59). Atribacterial ORFs encoding the Rnf complex were expressed at the majority of depths (Fig. 4B) indicating this is indeed an important mechanism of anaerobic ATP synthesis.

We also detected active transcription of pyruvate-formate lyase and acetate kinase, and formate dehydrogenase which is reversible (63), and therefore the WLP could be used for either catabolic or anabolic purposes (Fig. 5). However, the expression of an H_4_F: B_12_ methyltransferase from “*Ca*. Atribacteria” is a direct indication that the WLP is functioning as an acetogenic pathway as opposed to methanogenesis, because acetogens use 5-methyltetrahydrofolate: corrinoid (H_4_F: B_12_) methyltransferase, whereas methanogens use tetrahydromethanopterin methyltransferase in the last step of methyl synthesis prior to acetyl synthesis (66). Expression of the H_4_F: B_12_ methyltransferase from “*Ca*. Atribacteria” was relatively high in the deeper samples at 8.2 and 9.7 mbsf (Fig. 4B) where the steadily increasing abundance of 16S rRNA genes with depth reached their peak values (Fig. 1B).

“*Ca*. Atribacteria” expression of transporter-encoding genes for multiple types of sugar (i.e. hexose, hexulose, pentose, pentulose, ribose) and glycosidases (Fig. 4B) indicates a potential preference for sugar-based substrates. Sugars are in anoxic sediments as an energy substrate from bacterial necromass (e.g., ribose containing nucleic acids DNA and RNA) and could be used as a fermentation substrate (67). The only detectable gene expression from “*Ca*. Atribacteria” observed at 15.9 m depth indicates sugar transport, peptidases, fructose biphosphate and phosphogluconate aldolase activities (Fig. 4B), again pointing that utilization of necromass (sugars and peptides from dead cells) is an important activity for long-term survival.

Fermentation products and other oxidized organic substrates (i.e. alcohols, ketones, aldehydes, carboxylic acids) become toxic if they accumulate in the cell (68, 69). The gene expression from “*Ca*. Atribacteria” shows a metabolic potential for aldehydes to be re-oxidized in secondary fermentations via a bacterial micro-compartment (BMC), regenerating additional reduced ferredoxin and NADH in the process (70, 71). We obtained three contigs from ‘Ca. Atribacteria’ with ORFs encoding BMC shell proteins syntenous with Rnf, aldehyde dehydrogenase, and 2-deoxy-D-ribose 5-phosphate aldolase (DERA) (Fig. 4C), similar to the already reported genome assembled from “*Ca*. Atribacteria” (57). These genes were co-expressed at multiple depths (Fig. 4B), suggesting that the BMC and Rnf-based establishment of the Na^+^ ion gradient for ATP production are connected (Fig. 5). Collectively, these findings indicate that “*Ca*. Atribacteria” may use a BMC to recycle toxic intermediates produced during secondary fermentations for additional energy (Figs. 4-5). The ability to use a BMC to sustain secondary fermentations, in addition to primary fermentation of sugars (Figs. 4-5), may optimize energy conservation (e.g., increase ATP yields per mole substrate oxidized) under the extreme energy limiting subseafloor conditions. Similar to our results, a dominance of “*Ca*. Atribacteria” in organic lean sediments has been observed previously during IODP expedition 313 to the New Jersey shelf (27). However, we acknowledge that biases inherent to all methods for determining the abundance of Bacteria and Archaea make quantitative comparisons difficult (5), and thus interpret the qPCR data showing a relatively lower abundance of Archaea (Fig 1B) with caution.

Although fermenters and acetogens are often outcompeted by SRB in organic-rich subseafloor sediments, the poor fitness of SRB in the sampled environment (Figs. 1B-C), compared to “*Ca*. Atribacteria”, may be related to the relatively low SO_4_^2−^ reduction rates (Fig. 1A) and relatively low rates of organic matter burial at our sampled site (11, 19). In contrast, Chloroflexi related to Dehalococcoidia, which are predicted (homo)acetogens with metabolic potential to degrade complex organic compounds (72, 73), are relatively abundant in these deep-sea anoxic clay (Figs. 1B-C) and transcribe genes involved in cell division (Figs. 3B and S9). Our results lead us to speculate that organic-lean, anoxic abyssal subseafloor sediment thus represents a niche that favors the reproduction of Chloroflexi and “*Ca*. Atribacteria” over other microbes including SRB. Moreover, in culture experiments from the first cultivated representative of the Atribacteria, the presence of an H_2_-consuming methanogenic partner increased the growth rate of Atribacteria by more than 100 fold (43). Based on this experimental result, we speculate that the relatively high abundance of the Atribacteria in our sampled ancient anoxic sediment could be attributed to an as-of-yet unidentified syntrophic or semi-syntrophic partner organism, from the Chloroflexi, for example Dehalococcoidia (74, 75).

Our data show that reproduction can occur in the subseafloor over multimillion year timescales. Although RNA from the “*Ca*. Atribacteria” was no longer detectable below 16 mbsf, metagenomes and 16S sequencing show that they subsist and remain the dominant group down to at least 29 mbsf. The extractable DNA with our protocol does not target extracellular DNA (20), and thus the DNA in these deeper sediments likely does not derive from “dead” DNA preserved from once living cells. Therefore, we assume that “*Ca*. Atribacteria” remain viable, and potentially active, in the deeper sediments but at cell abundances and transcriptional activities that are too low for our current RNA-based methods. Thus the survival of the “*Ca*. Atribacteria”, associated with gene expression at relatively low levels, supports the hypothesis that reduced metabolic activity is a fitness advantage (16) in the energy-starved subseafloor.

## Materials and Methods

### Sampling

All samples were taken by Cruise KN223 of the R/V Knorr in the North Atlantic, from 26 October to 3 December 2014 (Woods Hole, MA to Woods Hole, MA). At site KN223-15 (33°29.0’ N, 54°10.0’ W, water depth 5515 m), successively longer sediment cores were retrieved using a multicorer (~0.4 m), gravity corer (~3 m) and the Woods Hole Oceanographic Institution (WHOI) piston-coring device (~28 m). Additional details of sampling are published (11). Dissolved oxygen concentrations in the core sections (Fig. S1) were measured with optical O_2_ sensors as described previously (76). Concentrations of dissolved SO_4_^2−^ were measured as published previously (6), and SO_4_^2−^ reduction rates were calculated as previously described (14). Sediment subcores were retrieved on the ship aseptically using end-cut sterile syringes and kept frozen at −80 °C until extraction in the home laboratory.

### DNA extraction, quantitative PCR and 16S rRNA gene libraries

For each sampled depth, we performed four biological replicates of DNA extraction. Total DNA was extracted from 0.7 g of sediment as previously described (20). DNA templates were diluted 10 times in ultrapure PCR water (Roche) and used in qPCR amplifications with updated 16S rRNA gene primer pair 515F (5′-GTG YCA GCM GCC GCG GTA A -3′) with 806R (5′-GGA CTA CNV GGG TWT CTA AT -3′) to increase our coverage of Archaea and marine clades (Parada et al., 2016) and run as previously described (38). All qPCR reactions were set up in 20 μL volumes with 4 μL of DNA template, 20 μL SsoAdvanced SYBR Green Supermix (BioRad), 4.8 μL Nuclease-free H2O (Roche), 0.4 μL primers (10 μM; biomers.net) and 0.4 μL MgCl2 and carried out on a CFX-Connect qPCR machine for gene quantification. For 16S rRNA genes, we ran 40 PCR cycles of two steps corresponding to denaturation at 95 °C for 15 sec, annealing and extension at 55 °C for 30 sec. To measure the abundance of dsrB genes, we used a previously described assay (77) with the primer pair dsrB4-R (5′-GTG TAG CAG TTA CCG CA -3′) with dsrB2060F (5′-CAA CAT CGT YCA YAC CCA GGG -3′). All qPCR reactions were set up in 20 μL volumes with 4 μL of DNA template and performed as previously described (78). Gel purified amplicons of the 16S rRNA, dsrB and mcrA genes were quantified in triplicate using QuantiT dsDNA reagent (Life Technologies) and used as a standard. An EpMotion 5070 automated liquid handler (Eppendorf) was used to set up all qPCR reactions and prepare the standard curve dilution series spanning from 10^7^ to 10^1^ gene copies. Reaction efficiency values in all qPCR assays were between 90 % and 110 % with R^2^ values >0.95 % for the standards.

For 16S rRNA gene library preparation, qPCR runs were performed with barcoded primer pair 515F and 806R. All 16S rRNA gene amplicons were purified from 1.5 % agarose gels using the QIAquick Gel Extraction Kit (Qiagen), quantified with the Qubit dsDNA HS Assay Kit (Thermo Fisher Scientific), normalized to 1 nM solutions and pooled (Pichler et al., 2018). Library preparation was carried out according to the MiniSeq System Denature and Dilute Libraries Guide (Illumina). Sequencing was performed on all four biological replicates (Fig. S2) on the Illumina MiniSeq platform at the Geo-Bio LMU Center. We used USEARCH version 10.0.240 for MiniSeq read trimming and assembly, OTU picking and 97 % sequence identity clustering (79), which we showed previously captures an accurate diversity represented within mock communities sequenced on the same platform (38). OTU representative sequences were identified by BLASTn searches against SILVA database version 132 (80). To identify contaminants, 16S rRNA genes from extraction blanks and dust samples from the lab were also sequenced in triplicate (38). These 16S rRNA gene sequences were used to identify any contaminating bacteria (e.g. *Acinetobater*, *Bacillus*, *Staphylococcus*) and selectively curate the OTU table of our anoxic abyssal clay samples prior to downstream analysis. The 30 most abundant OTUs were aligned with SINA online v.1.2.11 (81) and plotted in a Maximum Likelihood RAxML phylogenetic tree (82) (Fig. S3), against the SILVA 16S rRNA SSU NR99 reference database version 132 (80) using ARB (83). Closest environmental sequences with nearly full-length sequences (>1400 bp) were selected as taxonomic references and used to calculate trees implemented with the bacterial and archaeal filter and advanced bootstrap refinement selecting the best tree among 300 replicates (83). Partial OTU sequences were added to the tree using the maximum parsimony algorithm without allowing changes of tree topology (Figs. 2 and S3).

### Metagenome libraries

Whole genome amplifications were performed on DNA extracts at dilution 10 times through a multiple displacement amplification (MDA) step of 6 to 7 hours, using the REPLI-g Midi Kit (QIAGEN) and following the manufacturer’s instructions. We added SYBR green I (Invitrogen) at 1000 × concentration to visualize the incubation on the CFX-Connect qPCR machine with a picture taken every 10 min. Amplification was stopped after reaching the exponential increase by heating to 65 °C for 3 min. Metagenomic libraries were prepared using the Nextera XT DNA Library Prep Kit (Illumina), then quantified on an Agilent 2100 Bioanalyzer System (Agilent Genomics) and normalized with the Select-a-Size DNA Clean and Concentrator MagBead Kit (Zymo Research) as previously described (20).

### RNA extractions and metatranscriptome libraries

Total RNA extractions were obtained following a previously published protocol (84). In brief, RNA was extracted from 3 g of sediments using the FastRNA Pro Soil-Direct Kit (MP Biomedicals) following the manufacturer’s instructions, with the addition of 4 μL glycogen (0.1 g × mL^−1^) to increase yield during precipitation of the RNA pellet, and final elution in 40 μL PCR-grade water (Roche). Extraction blanks were processed alongside to assess laboratory contamination. RNA extracts were quantified using the QuBit RNA HS Assay Kit (Thermo Fisher Scientific). DNAse treatment, synthesis of complementary DNA and library construction were processed on the same day from 10 μL of RNA templates using the Trio RNA-Seq kit protocol (NuGEN Technologies). Libraries were quantified as described above. All libraries were diluted to 1 nM and pooled for further sequencing on the MiniSeq platform (Illumina).

### Gene identification and normalization in metagenomes

The SqueezeMeta (85) metagenomic analysis pipeline was used for downstream analysis of metagenomic reads in co-assembly mode. For adapter removing, trimming and quality filtering, we used Trimmomatic set to: leading = 8; trailing = 8; sliding window = 10: 15; and minimum length = 30 (86). Contigs were assembled to minimum length of 200 bp using Megahit assembler (87). ORFs for genes and rRNAs were called using Prodigal (88), rRNAs genes were determined by barrnap (89). RDP classifier were used for the classification of 16s rRNA genes (90). Diamond software (91) was deployed for taxonomic assignment of retrieved gene homologies against the Genbank, eggNOG v. 4.5 (92) and KEGG databases (93). Cut-off values for assigning hits to specific taxa were performed at e-value 1 × e^−3^, minimum amino acid similarity of 40 for taxa and 30 for functional assignment, using SqueezeMeta with default settings. Reads were mapped onto contigs and genes, using Bowtie2 (94). Coverage and transcripts per million (TPM) values were calculated using SqueezeMeta. For binning, we used Maxbin 2.0 (95) and MetaBAT (96), and bins generated by the two different algorithms were merged into one single dataset using DAS Tool (97). Bin completeness and contamination were checked using CheckM (98). Further analysis of metagenome-assembled genomes (MAGs) based on results from SqueezeMeta was achieved using Anvi’o v. 6.2 (32) and MAGs selected by DAS tool further refined manually based on hierarchical clustering of contigs. The complete workflow using DAS Tool separates 14 bins whose taxonomic affiliations are uncertain. Manual curation of these results points to Aerophobetes, Chloroflexi and Atribacteria as potential bin affiliations (Table S2). In comparison, from the manual curation of Maxbin results, we produced 31 bins (17-59%) among which 12 bins are clearly assigned to “*Ca*. Atribacteria” with different levels of genome completeness (Fig. S4).

Because the MDA step produced many short fragments that did not allow high-quality binning and full genome completion, assigning taxonomic affiliation to metagenomic and metatranscriptomic data is challenging (99). For annotating putative functions of ORFs in metagenomes and metatranscriptomes from particular “higher-level” taxonomic groups of microorganisms (34, 35), we also applied a bioinformatics pipeline involving a large aggregated genome database of predicted proteins including the SEED (www.theseed.org) and NCBI RefSeq databases updated with all predicted proteins from recently described high-quality draft subsurface MAGs and single-cell assembled genomes (SAGs) from the NCBI protein database and all fungal genomes from the NCBI RefSeq database (34, 84). The total number of predicted proteins in the updated database was 37.8 million. This approach assigns ORFs to higher-level taxonomic groups (35). As is the case in all metagenomic studies, the incomplete nature of genomes in databases, together with the lower representation of sequenced genomes from candidate clades compared to cultured ones, make it likely that our pipeline misses annotation of ORFs that are derived from as-of-yet unsequenced Atribacteria genomes. We acknowledge that some genes in databases annotated as being present in Atribacteria might have been assigned to bins according to criteria that differ from study to study (Fig. S7).

### Gene identification and normalization in metatranscriptomes

Paired-end reads were trimmed and assembled into contigs using CLC Genomics Workbench 9.5.4 (https://www.qiagenbioinformatics.com/), using a word size of 20, bubble size of 50, and a minimum contig length of 300 nucleotides. Reads were then mapped to the contigs using the following parameters (mismatch penalty = 3, insertion penalty = 3, deletion penalty = 3, minimum alignment length = 50% of read length, minimum percent identity = 95%). Coverage values were obtained from the number of reads mapped to a contig divided by its length (i.e. average coverage). Only contigs with an average coverage >5 were selected for ORF searches, and downstream analysis (20, 34, 84). This protocol does not assemble ribosomal RNA (rRNA), and thus results are only discussed in terms of messenger RNA (mRNA). We then performed even further stringency controls by removing any contig that had less than 5 × coverage, e.g. reads per kilobase mapped (RPKM). The final resulting dataset of contigs was then used for ORF searches and BLAST analysis. Protein encoding genes and ORFs were extracted using FragGeneScan v. 1.30 (100) and functionally annotated against an large aggregated genome database (20) (34, 35) containing predicted proteins from all protist, fungal, bacterial, and archaeal genomes and MAGs in the JGI and NCBI databases using DIAMOND version 0.9.24 (91). This database, which we refer to as ‘MetaProt’ also contained all ORFs from all of the transcriptomes of microbial eukaryotes from the MMETS project (101), and we removed any hits to photosynthetic eukaryotic algae as contaminants. This custom MetaProt database that we used for this study is available as a single 32GB amino acid fasta file on the LMU Open Data website (doi.org/10.5282/ubm/data.183). Cut-off values for assigning hits to specific taxa were performed at a minimum bit score of 50, minimum amino acid similarity of 60, and an alignment length of 50 residues. All scripts and code used to produce the analysis have been posted on GitHub (github.com/williamorsi/MetaProt-database), and we provide a link to the MetaProt on the GitHub page, as well as instructions within the scripts regarding how to conduct the workflows that we used.

For the metatranscriptomes, normalization of the relative abundance of ORFs was based on the number of unique ORFs assigned to a group (e.g., “*Ca.* Atribacteria”) a fractional percentage of total ORFs detected. We normalized expression in this manner, as opposed to more conventional procedures such as RPKM because we found that the RNAseq kit we used has an amplification step (SPIA amplification, Trio RNAseq Ovation kit, NuGen) that biases the relative abundance of reads mapping to contigs when normalized using RPKM. For example, the RPKM value for the same ORF across technical replicates was found to have very large (orders of magnitude) variability in RPKM. In contrast, the total number of unique ORFs (e.g., presence absence of an expressed ORF) assign to specific groups (e.g., “*Ca*. Atribacteria”) was highly consistent between technical replicates. We assume that this technical variation in the RNAseq data is associated with randomized SPIA amplification of different transcripts, and or fluctuations in the number of mRNA molecules in technical replicate tubes due to the highly labile nature of RNA during the extraction and library prep procedure. For this reason, we normalized the relative abundance of ORFs assigned to a specific group based on presence/absence of expressed ORFs which was highly consistent between technical replicates despite the SPIA amplification. If significantly higher numbers of unique ORFs are detected from a particular group compared to other groups it can be attributed to a relatively higher transcriptional activity.

COG categories were assigned by searching the ORFs against the COG database (102) using BLASTp. Metagenomic raw reads and metatranscriptomic atribacterial contigs were mapped with high stringency against a previously sequenced subseafloor MAG of “*Ca*. Atribacteria” that has relatively high completeness (88%) as a reference (103), using Geneious 8.1.9. The average coverage after mapping against GenBank references no. NCRO00000000.1 to .700 (103) was 224 (± 131) and 93 % of ORFs in the reference MAG were detected in our subseafloor metagenomes and metatranscriptomes (Fig. S10). Consensus sequences were exported and annotated using the online tool RAST v. 2.0 (104). Taxonomic assignment of protein-encoding genes to “*Ca*. Atribacteria” clade JS1 was further confirmed in our metagenomes (Fig. S6) by selecting and aligning 31 phylogenomic markers from the corresponding metagenomic ORFs and 36 atribacterial reference genomes obtained from the NCBI database, using AMPHORA2 (33). Statistical analyses of beta-diversity were performed using RStudio v. 3.3.3 with the Bioconductor package (105). Contig rings (Fig. S10) were plotted using Circos v. 0.69-6 (106).

Data are publicly available through NCBI BioProject PRJNA590088. Metagenomes and metatranscriptomes have accession number SAMN13317858 to SAMN13317880. The 16S data are available in SRA BioSample accessions SAMN10929403 to SAMN10929517 and SAMN13324854 to SAMN13324920. Additional data related to this paper may be requested from the authors.

## Acknowledgements

This work was supported primarily by the Deutsche Forschungsgemeinschaft (DFG) project OR 417/1-1 granted to W.D.O. Preliminary work was supported by the Center for Dark Energy Biosphere Investigations project OCE-0939564 also granted to W.D.O. The expedition was funded by the US National Science Foundation through grant NSF-OCE-1433150 to S.D., and R.P. R.W.M. led the expedition. Shipboard microbiology efforts were supported by the Center for Dark Energy Biosphere Investigations (C-DEBI grant NSF-OCE-0939564). This is a contribution of the Deep Carbon Observatory (DCO).

## Author contributions

W.D.O. conceived the work and experimental approach. A.V., W.D.O., Ö.K.C., and S.V. contributed to the laboratory/bioinformatics analyses and experimental work. R.W.M., D.C.S. and R.P. obtained the samples during the KN223 R/V Knorr oceanographic expedition. W.D.O., R.W.M., D.C.S., A.V., S.V., and S.D. discussed and wrote the manuscript and commented on the paper.

